# Restoration of a pond: monitoring water quality and macroinvertebrate community succession

**DOI:** 10.1101/2021.10.19.465034

**Authors:** Abigail Kuranchie, Aaron Harmer, Barbara Evans, Dianne H. Brunton

## Abstract

To determine the success of restoration programmes, knowledge of the temporal dynamics in community structure and processes is vital. The water quality and macroinvertebrate community structures of a newly created and an established pond within the same ecosystem were sampled bi-monthly over a year to monitor the development of the new pond. The water quality measures of the ponds were significantly different. Conductivity, salinity, and total dissolved solids levels were also different between the ponds. The colonisation of a macroinvertebrate community in the new pond was rapid, resulting in a 75% resemblance to the established pond by the end of the first year. The pond was colonised by non-insect taxa like Crustacea and Gastropod and then by insects. There was a significant difference in the macroinvertebrate communities of the ponds due to temporal taxonomic composition differences. The high abundance of *Diplacodes* spp. (perchers), *Physa* spp. (left-handed pond snail), and Ostracod (seed shrimp) in the new pond contributed to the difference in the community between the two ponds. Cladocera (water fleas) dominated the macroinvertebrate community, and the highest abundance was recorded in August for both ponds. Our results suggest that a newly created pond can have a comparable macroinvertebrate community to nearby established ponds within a year.

## Introduction

The construction of new ponds and the restoration of freshwater ecosystems to compensate for losses of these systems have become necessary in recent times (Coccia et al. 2016). Assessing the role and value of these restored or new ecosystems to determine if they complement existing systems or replace the natural systems is vital for conservation and management. However, to achieve a restoration goal, knowledge of the temporal dynamics in community structure is essential. Monitoring provides essential benchmarks for restoration programmes and planning, supporting projects to make realistic milestones for measuring success (Burbage 2005).

Ponds have economic, cultural, social and conservation values. They help maintain the water table and are home to diverse organisms, including algae, macroinvertebrates, amphibians, and fish. Artificial ponds are useful ecosystems for studying the successional process in freshwater ecosystems. They allow for studying the species composition from the onset of creating the habitat until the community stabilises (Olmo et al. 2016). For organisms to colonise and be established successfully in a new habitat, for example, after a natural disaster has removed existing ecosystems, they must actively or passively disperse to the site. Coloniser organisms must be able to establish successfully in the presence of predators, parasites, and competitors. Finally, the habitat must be suitable for survival and reproduction (Wood et al. 2003; Olson et al. 2016).

Nonetheless, in situations where there is a relic of biodiversity from sources such as egg or seed banks, colonisation is possible when conducive environmental conditions occur (Bilton et al. 2001). Species composition changes throughout this succession process until a stable community is attained (Ruhí et al. 2013). Therefore, information on the temporal dynamics is essential in improving the knowledge of succession and providing information to construct similar ecosystems (Miguel Chinchilla et al. 2014).

Macroinvertebrates are one of the first organisms to colonise aquatic habitats. International studies on succession have found that macroinvertebrate communities become fully established, reaching high densities within the first year of pond creation (Solimini et al. 2003; Cañedo-Argüelles and Rieradevall 2011; Ruhí et al. 2013). Taxa with high dispersal abilities are early colonisers, followed by passive dispersers or non-flying taxa (Cañedo-Argüelles and Rieradevall 2011). Macroinvertebrates are good models to study succession and biological restoration as they are primary consumers and play a vital role in the energy exchange in the ecosystem. Macroinvertebrates are also useful in monitoring programmes because they are sensitive to changes in ecological processes like primary productivity and nutrient and energy flow (Coccia et al. 2016). Newly created ponds show an increase in macroinvertebrate diversity over time due to the presence of many unoccupied niches until they reach carrying capacity (Coccia et al. 2016). However, most newly created ponds lack macrophytes that create additional habitats for macroinvertebrates (Coccia et al. 2016).

Studies on colonisation/succession of aquatic ecosystems have been carried out in numerous countries, including the United States of America (Layton and Voshell Jr 1991; Batzer DP and Resh 1992), in Europe (Solimini et al. 2003; Boix et al. 2004; Culioli et al. 2006; Cañedo-Argüelles and Rieradevall 2011), in Africa (Guiral et al. 1994) and Australia (Bayly 2001). There is, however, little information on the processes of primary succession of freshwater macroinvertebrates in artificially restored ponds in New Zealand.

Newly created ponds provide an ideal system for studying succession because the physical, chemical, and biological community changes can be easily monitored (Ruhí et al. 2011). The construction of a new pond in the Matuku Link provided a unique opportunity to study primary succession in a natural system. This study took advantage of a unique opportunity to assess pond restoration from formation to full ecological function and compare it to an established pond within the same landscape. The aims were to 1) monitor and compare the water quality trends of both the established and the new pond, 2) monitor and compare the macroinvertebrate community structure, composition, and diversity of both the new and established ponds; and 3) describe the succession of macroinvertebrates in the new pond.

## Methods and methods

### Description of study ponds

The established pond, also known as the Dr John pond (GPS coordinates; Latitude: −36.865, longitude: 174.489), is artificial, shaped like a speech bubble and covers 147m^2^ and is about 0.5m deep when full. The pond is surrounded by predominantly native vegetation and is connected to a marshland on its eastern edge. This riparian vegetation shades up to 60 % of the pond surface, but this depends on the time of day and time of year. The pond is dominated by *Myriophyllum aquaticum* (parrot feather), an introduced submerged aquatic plant.

The 180 m^2^ oval-shaped new pond is also artificial (Latitude: −36.864, Longitude: 174.489) and leads to a swamp at the west end. This pond was constructed to serve as additional freshwater habitat to enhance the Matuku Link reserve’s conservation function. It was dug to a depth of 2m, but due to rapid siltation, the average depth recorded during this study was 1.2m. The riparian vegetation is dominated by *Ranunculus* spp. (buttercups). Due to a slope on one side of the pond, it is expected to receive runoff from the adjacent walking path.

Water quality was measured, and macroinvertebrates sampled from the ponds bi-monthly from February 2019 to December 2019. The measured water quality parameters included pH, temperature, total dissolved solids (TDS), percentage dissolved oxygen (% DO), salinity, ammoniacal nitrogen (NH_3_-H), orthophosphate (PO_4_^3-^), and nitrate (NO_3_-N). Water quality testing was done on-site using a calibrated Hanna multiparameter probe (Model H198194) (Waqas et al. 2018) and the nutrients concentrations with HACH nutrient test kits. Algal biomass in the pond was measured using the spectrophotometry method. Macroinvertebrates were sampled with a D-frame net simultaneously as the water quality testing was carried out. Sampling was done for a minute each at three different spots around each pond. Macroinvertebrate samples were sorted and identified to the family or genus level for insects and molluscs using identification keys. Earthworms, Nematodes, seed shrimps and hydra were counted but not identified further.

#### Data analyses

Water quality parameters were transformed and then normalised for further analyses based on the assumptions of statistical testing. A paired Student t-test was used to test for significant differences between the overall average value of the individual physicochemical parameters measured in the two ponds. The difference in each pond’s physicochemical composition was tested using permutational multivariate analyses of variance (PERMANOVA) (Clarke and Warwick 2001). Similarly, differences in macroinvertebrate composition between the ponds were documented using Permanova based on a Bray Curtis similarity calculated on the transformed (fourth root) and standardised data (Bray and Curtis 1957). The similarity of percentages (SIMPER) analysis was used to determine the taxa that contributed to the macroinvertebrate communities’ differences. Various diversity indices such as Margalef’s richness, Pilou’s evenness, Shannon-Wiener diversity, and Average taxonomic distinctness were used to analyse different aspects of each pond’s biodiversity. All diversity indices were computed using PRIMER-E (version 7) (Clarke and Gorley 2006).

## Results

### Physicochemical water quality parameters

The new pond recorded higher levels of conductivity, salinity, and total dissolved solids. Additionally, there was more variability in the temporal trend of the parameters measured in the new pond. The average temperature in the new pond (17.05 °C ± 5.7 °C) was higher and was more variable than the established pond (15.67 °C ± 5.03 °C) (Table 1). Conductivity was higher during the colder months than the warmer months for both ponds, but there was higher variability in the conductivity levels in the new pond. The pH ranged from weak acidity to weak base (5.53–7.80) throughout the study period in both ponds. The pH in the established pond ranged between 7.8 and 6.13, and in the new pond, between 7.21 and 5.53. Percentage dissolved oxygen concentration (% DO) in both ponds fluctuated throughout the year. The average ammonia concentration in the new pond (0.038 ± 0.04mgl^-1^) was slightly higher than that of the established pond (0.03 ± 0.03mgl^-1^). Chlorophyll ‘a’ concentration in the new pond was highest in February (4.4 x 10^-2^ mgm^-2^) and lowest in October (4.3×10^-4^ mgm^-2^). In the established pond, the chlorophyll ‘a’ concentration was highest in February (0.2mgm^-2^) but lowest in December (5.2×10^-4^ mgm^-2^). There was more variability in the temporal trend of the physicochemical water quality parameters measured in the new pond than in the established pond. The water quality in the established pond in October and December were most similar; however, the quality in February and June was most dissimilar, as shown in the non-multidimensional scaling (nMDS) plot (Figure 1a). The high abundance of *Diplacodes* spp., *Physa* spp., and Ostracod in the new pond contributed to the difference in the community between the ponds.

**Table 1:** A summary of physicochemical water quality parameters and macroinvertebrate diversity measures of the new and established pond. The table shows the means ± standard deviations (SD) and the range of values. The parameters have their units in brackets. Significant p-values at 95% confidence from a paired t-test are in bold.

| Parameters | New Pond (n = 6) |  | Established Pond (n = 6) |  | t | p |
| --- | --- | --- | --- | --- | --- | --- |
| Physicochemical parameters | Mean ± SD | Range | Mean ± SD | Range | | |
| pH | 6.77 ± 0.65 | 5.53–7.25 | 6.79 ± 0.55 | 6.13–7.80 | -0.12 | 0.9 |
| % DO | 34.71 ± 32.41 | 2.7–92.9 | 30.84 ± 22.04 | 13.86–73.43 | 0.56 | 0.6 |
| Conductivity ($\mu\text{scm}^{-1}$) | 240.4 ± 32.2 | 207–285.7 | 214.7 ± 25.3 | 181–252 | 3.93 | **0.01** |
| TDS (ppm) | 120.3 ± 16.0 | 104–142.7 | 107.3 ± 12.9 | 90.7–127 | 3.90 | **0.01** |
| Salinity (psu) | 0.11 ± 0.01 | 0.1–0.14 | 0.10 ± 0.01 | 0.09–0.12 | 3.16 | **0.02** |
| Temperature ($^{\circ}\text{C}$) | 17.05 ± 5.77 | 10.62–24.65 | 15.67 ± 5.03 | 9.86–22.65 | 1.34 | 0.2 |
| NO<sub>3</sub>-N ($\text{mg l}^{-1}$) | 0.29 ± 0.21 | 0.02–0.49 | 0.32 ± 0.24 | 0.01–0.51 | -0.30 | 0.1 |
| PO<sub>4</sub><sup>3-</sup> ($\text{mg l}^{-1}$) | 1.25 ± 0.98 | 0.24–2.36 | 1.98 ± 0.94 | 0.09–2.68 | -1.72 | 1 |
| NH<sub>3</sub>-H ($\text{mg l}^{-1}$) | 0.03 ± 0.04 | 0.01–0.12 | 0.03 ± 0.03 | 0.01–0.10 | <-0.02 | 0.4 |
| Chlorophyll 'a' $\times 10^{-3} \text{mgm}^{-2}$ | 9.7 ± 17.0 | 0.4–43.8 | 38.3 ± 90.7 | 0.5–223.5 | -0.94 | 0.4 |
| Macroinvertebrate's diversity measures |  |  |  |  |  |  |
| Margalef's richness 'D' | 2.7 ± 0.3 | 2.2–3.1 | 2.7 ± 0.4 | 2.2–3.13 | -0.19 | 0.9 |
| Pilou's evenness 'J' | 0.8 ± 0.1 | 0.6–0.8 | 0.7 ± 0.1 | 0.5–0.9 | 1.4 | 0.2 |
| Shannon-Wiener diversity 'H' | 2.17 ± 0.14 | 1.9–2.3 | 0.8 ± 0.4 | 1.3–2.5 | 1.4 | 0.2 |
| Average taxonomic distinctness '$\Delta^{+}$' | 82.84 ± 3.42 | 77.9–88.2 | 81.3 ± 3.7 | 77.6–88.2 | **3.17** | **0.02** |

**Figure 1:**
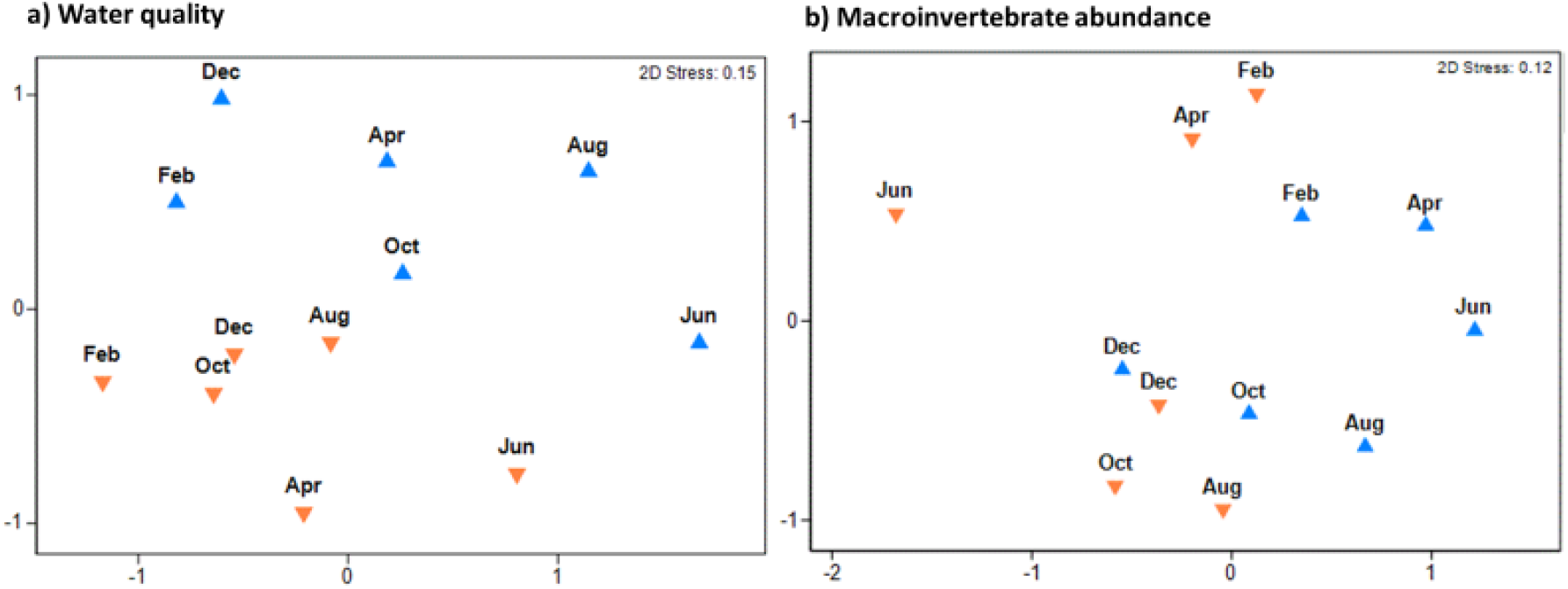
An nMDS plot of; a) the water quality of the established and new ponds based on Euclidian distance measure of transformed and normalised physicochemical water quality data, and b) macroinvertebrates abundance (over time) based on Bray Curtis similarity measure on a fourth root transformed data. Blue triangles indicate the new pond data points, and orange triangles indicate data points in the established pond.

### Macroinvertebrate community structure

A total of 7448 macroinvertebrates were sampled from the two ponds from February to December 2019. Forty-two taxa were recorded in both ponds, and 16 were consistently present in 50% of the samples. The taxa Oligochaeta and *Cura* spp. were present in all the samples. Table 2 summarises the occurrence of macroinvertebrates in the two ponds. The minimum number of taxa occurred in June from both ponds, and the maximum was in August for the new pond (23) and October for the established pond (22). Thirteen taxa were recorded from both ponds during the first sampling period in February. Although the number of taxa increased marginally in the new pond during April and June, the established pond had 13 taxa recorded for April and June. In the established pond, non-insect taxa dominated the macroinvertebrate community. In the new pond, non-insect taxa dominated the community structure for all months sampled except June (Table 2).

**Table 2:** A comparison of the macroinvertebrate composition in the new and established ponds sampled from sampled bi-monthly from February to November.

| Descriptive measures | New pond | Established pond |
| --- | --- | --- |
| Percentage of overall macroinvertebrates sampled | 55% | 45% |
| Average individuals per sample | 688 | 554 |
| The average number of taxa per sampling session | 17 | 16 |
| Total number of taxa present | 36 | 35 |
| Five most common taxa | Cladocera, Ostracod, Orthocladinae, Cyclopoida and Hydrachinidae | Oligochaeta, *Cura*, Chironomus, Calanoida and Hydrachinidae |
| Rare and exclusive taxa | Stratiomyidae, Pelecorrhynchidae, and Stictocladius | Dolichopodid, Mesovilia, Shaeriidae and Paraleptamphopus |
| Percentage of exclusive taxa | 14% | 16% |
| Relative abundance of insects' taxa | Lowest in August, highest in June | Lowest in April, highest in December |

#### Univariate Diversity Measures

The macroinvertebrates in the new ponds were more diverse. The average Shannon-Wiener diversity indices were 1.9 ± 0.4 and 2.1 ± 0.1 for the established and new pond, respectively. The highest Shannon Weiner diversity in the new pond was observed in October (2.3) and the lowest in August (1.9). The highest diversity was observed in the established pond in December (2.5) and the lowest in April (1.3).

Macroinvertebrates were more evenly distributed in the new pond than the established pond. The average Pilou’s evenness was 0.70 ± 0.13 and 0.77 ± 0.08 for the established and new pond, respectively. The highest evenness was recorded in December for both ponds. The macroinvertebrate richness was higher in the established pond. The average Margalef’s richness for the established and new pond were 2.7 ± 0.3 and 2.6 ± 0.4, respectively (Table 1). Macroinvertebrate richness in the established pond was highest in June (3.1) and lowest in April (2.2). In the new pond, the highest richness (3.0) was observed in August and the lowest (2.1) in December. Macroinvertebrates sampled in April were the least taxonomically related in the ponds, while the macroinvertebrates sampled in August were most taxonomically related.

#### Temporal dynamics of macroinvertebrates in the ponds

Macroinvertebrate composition was most similar (75.2%) between the ponds in December and the least similar (31.9%) in June. Additionally, there was more variability in the macroinvertebrate community between sampling sessions in the established pond. There was a significant difference (F _1, 5_ = 2.22, p = 0.02) between the community composition in the two ponds, as shown in the nMDS plot (Figure 1b).

## Discussion

### Water quality trends

This study is one of the few New Zealand studies to assess the rapid changes in a newly established pond using macroinvertebrate community and water quality parameters. The study monitored the water quality and macroinvertebrate community between a newly created pond and a nearby, comparable established pond. The water quality in the ponds was different. The differences are likely caused by inter pond variations in conductivity, salinity and TDS, percentage of shade from overhanging trees, amount of pond surface area covered by vegetation (macrophyte) and pond depth (Angélibert et al. 2004). Macrophyte cover has been shown to substantially influence water quality in New Zealand lakes (Ministry for the Environment 2006). Though the established pond was slightly shallower and expected to be warmer, shade from trees appeared to have a higher effect on the water temperature than the slight difference in depth; hence they had similar temperatures.

### Macroinvertebrate trends and dynamics in the ponds

The development of macroinvertebrate structure and composition in the new pond was rapid, resulting in a community that was over 75% similar to the established pond within the one year of this study. Similar results from Spain have been reported by Olmo et al. (2016), where zooplankton community composition in a restored pond was similar to the nearby existing pond by the end of the first year. Although 42 macroinvertebrate taxa were recorded, few taxa dominated by abundance in both ponds. This result suggests that less abundant (rare) taxa use the ponds as a refuge habitat or are yet to establish thriving populations in the ponds. It might also be due to their life-history traits such as low mortality, high reproduction rate or adaptability to the pond’s prevailing conditions (Botwe et al. 2017). The low abundance of insect taxa may be because of their semi-aquatic life and seasonality in colonising habitats (Lahr et al. 1999; Suren and Lambert 2010; Olmo et al. 2016). Different taxonomic groups, especially zooplankton like rotifers, and ostracods, show variation in their peak abundance over time (Lahr et al. 1999), and this was also observed in this study. Cladocera populations primarily drove the high abundance of macroinvertebrates recorded in August (the peak of the winter season in New Zealand). The high abundance is associated with higher water volumes (due to high precipitation), which supports more diverse macroinvertebrates (Boix et al. 2000).

Measures of taxonomic distinctness reflect changes in community structure. The inconsistency in the taxonomic distinctness between the ponds over the study period can be attributed to water quality variables such as conductivity and TDS, impacting intra-pond habitat quality (Jeffries 2005). This irregularity may also indicate seasonality in the macroinvertebrate composition and community structure (Cañedo-Argüelles and Rieradevall 2011).

The significant differences in the macroinvertebrate community composition may be due to several factors, including taxa specific response to temporal changes in habitat qualities and physicochemical conditions in the ponds (Ruhí et al. 2011; Coccia et al. 2016). Different physiological needs of macroinvertebrate taxa may also explain the difference in community composition (Jeffries 2005). Additionally, the differences in the age, type and amount of macrophyte may contribute to differences in macroinvertebrate composition (Oertli et al. 2002; Jeffries 2005; Ruhí et al. 2011). Although some studies show a positive relationship between faunal richness and pond age (Miguel Chinchilla et al. 2014; Olmo et al. 2016), the results of our study did not; both ponds had similar taxa richness, similar to findings by Sun et al. (2019).

### Macroinvertebrate colonisation and succession in the new pond

We found the succession in the new pond went from domination by non-insect taxa such as Branchiopod, Clitellata, Gastropod and Ostracod to domination by insects by the 9^th^ month of its establishment (August) and non-insects in the proceeding months. A similar succession was reported by (Lahr et al. 1999; Coccia et al. 2016). The new pond was observed to have been colonised fast, with high diversity indices (evenness and richness) recorded for the first sampling period, just three months after pond construction. The fast rate of colonisation could be due to the proximity of possible sources of colonisers and the absence of geographical barriers between the new and the established pond (Cañedo-Argüelles and Rieradevall 2011). This result aligns with other studies showing that colonisation in new ponds can be rapid. This rapid colonisation has been reported in both experimental (Layton and Voshell Jr 1991; Batzer D and Boix 1999) and non-experimental studies globally (Scher and Thiery 2005; Cañedo-Argüelles and Rieradevall 2011; Ruhí et al. 2011; Taylor and Duggan 2012; Kim et al. 2014). Studies show that the most abundant taxa are crustaceans that dominate in the early succession stages of ponds and wetlands (Lahr et al. 1999), which is similar to our findings.

Additionally, the high average taxonomic distinctness and macroinvertebrate richness recorded during the first sampling period (February) indicates a fast colonisation rate compared to the study by Ruhí et al. (2011). The initial colonisation sequence in this study has been related to the dispersal abilities of the taxa Cladocera (Crustacea), Ostracod (Crustacea) and Odonata (Bilton et al. 2001; Cañedo-Argüelles and Rieradevall 2011; Olmo et al. 2016). There was a possibility of crustacean egg banks in the soil hatching when the pond was constructed, contributing to the observed result. Crustacean eggs survive prolonged periods of up to 20 years in dry conditions (Lahr et al. 1999; Bilton et al. 2001; Wissinger et al. 2009).

The relatively high biodiversity indices recorded for the new pond during the first sampling period could be due to the new niches available for colonisation after the pond’s construction (Cañedo-Argüelles and Rieradevall, 2011). Studies have shown that lentic ecosystems established close to established ones without geographical barriers have high macroinvertebrate composition within the first year of creation (Cañedo-Argüelles and Rieradevall 2011). Additionally, the relatively high abundance of a passive disperser, *Physa* spp. (Gastropod) in the first sampling could be due to the eggs’ transfer by other vectors such as waterfowl or individual dispersers (Cañedo-Argüelles and Rieradevall 2011; Taylor and Duggan 2012).

The new pond was established during the early summer (November 2018), which coincided with increasing temperatures and the beginning of high evaporation leading to lower water levels in the nearby established pond. Active fliers such as Odonata and obligate aquatic insects (Coleoptera and Hemiptera) might have found refuge in the new pond as the water level in the established pond lowered (Lahr et al. 1999; Olmo et al. 2016). Although successful colonisation and establishment depend on the colonisers’ dispersal ability, the habitat must be suitable for survival (Flory and Milner 2000; Olson et al. 2016).

The growth of emergent macrophytes by the second sampling period in April may have provided additional niches noted in the increase in Shannon-Wiener diversity and Margalef’s richness data (Flory and Milner 2000; Olmo et al. 2016). The fluctuations in the Shannon-Wiener diversity indices recorded in the new pond over the study period suggest that the new pond may still have some unoccupied niches (Coccia et al. 2016). These fluctuations may also be due to seasonality in the taxa, as observed in New Zealand lakes (Burns and Mitchell 1980). However, a long-term study may provide more insights into this. The major drawback of our study is the limited sample size and the lack of replication. This limitation can be resolved in an experimental approach by creating artificial ponds and monitoring their evolution over a more extended period.

## Conclusion

A difference in the ponds water quality was observed, and the physicochemical water quality variables were unstable. There was temporal variation in the water quality of both ponds. The new pond differed in environmental conditions by having less macrophyte cover and was deeper than the established pond. The macroinvertebrate community composition between the two ponds was also different; however, the biodiversity indices were similar. Non-insect taxa (Branchiopod, Clitellata, Gastropod and Ostracod) were the first to occur in high abundance in both ponds. The taxa’s dispersal ability probably determined the succession sequence observed in this study. Oligochaeta and *Cura* spp. were the common taxa in the ponds. However, the five most abundant taxa in the ponds were Chironomus, Orthocladinae, Hydrachinaedae, Oligochaeta, and Cladocera. Also, the established pond had more exclusive taxa than the new pond. Changes in the community composition and macroinvertebrates structure over the study period could be linked to environmental changes, including seasonal effects. The macroinvertebrate community composition in the ponds was 75 % similar at the end of the first year of the new ponds’ establishment. Therefore, the new pond provided additional and more diverse habitats for macroinvertebrates. It contributed to the overall increased biodiversity in the Matuku Link, which affirms that a cluster of ponds helps promote regional diversity.

## Author Contributions

Conceived and designed the experiment: AK, DHB, AH. Performed the experiment: AK, BE. Analysed data: AK, Wrote the paper: AK, DHB, BERestoration of a pond: monitoring water quality and macroinvertebrate community succession

Funding was provided by the Conservation Fund of Auckland Zoo and Massey University.

Acknowledgements, we thank the Matuku link Trust for permission to use the reserve for this study.

The authors declare that no competing interests exist.

## Data Availability Statement

All relevant data are within the paper and its Supporting Information files.

